# Genomic basis and phenotypic manifestation of (non-)parallel serpentine adaptation in *Arabidopsis arenosa*

**DOI:** 10.1101/2022.02.26.482078

**Authors:** Veronika Konečná, Marek Šustr, Doubravka Požárová, Martin Čertner, Anna Krejčová, Edita Tylová, Filip Kolář

## Abstract

Parallel evolution is common in nature and provides one of the most compelling examples of rapid environmental adaptation. In contrast to the recent burst of studies addressing genomic basis of parallel evolution, integrative studies linking genomic and phenotypic parallelism are scarce. Edaphic islands of toxic serpentine soils provide ideal systems for studying rapid parallel adaptation in plants, imposing strong, spatially replicated selection on recently diverged populations. We leveraged threefold independent serpentine adaptation of *Arabidopsis arenosa* and combined reciprocal transplants, ion uptake phenotyping, and available genome-wide polymorphisms to test if parallelism is manifested to a similar extent at both genomic and phenotypic levels. We found pervasive phenotypic parallelism in functional traits yet with varying magnitude of fitness differences that was congruent with neutral genetic differentiation between populations. Limited costs of serpentine adaptation suggest absence of soil-driven trade-offs. On the other hand, the genomic parallelism at the gene level was significant, although relatively minor. Therefore, the similarly modified phenotypes e.g., of ion uptake arose possibly by selection on different loci in similar functional pathways. In summary, we bring evidence for the important role of genetic redundancy in rapid adaptation involving traits with polygenic architecture.

## INTRODUCTION

Adaptation is a key evolutionary process allowing organisms to cope with changing environments. Identifying adaptation, however, is challenging at both phenotypic and genetic levels as multiple confounding signals such as evolutionary constraints, demographic or stochastic genetic processes may produce patterns resembling adaptation. Multiple populations facing the same environmental challenge may provide strong evidence for repeated adaptation within a species. If the same phenotypes independently emerge in similar habitats, it is more probable that these features evolved under natural selection than solely due to the stochastic forces (Lenormand et al., 2009). The same adaptive phenotypes can have a divergent genetic basis, e.g. when different genes within a pathway get under selection in different independent populations (e.g. Fang et al., 2021), or a similar genetic basis when adaptation sources from a shared pool of standing variation or from genes that are under a pleiotropic constraint (Hämälä and Savolainen, 2019). Despite numerous examples of parallel adaptation, we still have a limited understanding of the genetic basis of parallel phenotypic changes and their interplay with local environmental conditions.

The strength of correlation between environmental gradients and particular phenotypic traits, which are under selection in multiple adapted populations, provides an opportunity to quantify the extent of parallelism. However, independently evolved ‘adaptive’ phenotypes are rarely identical, representing rather a continuum of so-called non-parallel evolution (Stuart et al., 2017; Bolnick et al., 2018) in response to the complex interplay among the genetic basis of a trait, population history, and local environmental heterogeneity (Rosenblum et al., 2014; Fraïsse and Welch, 2019; James, Wilkinson, et al., 2021). Indeed, varying extent of phenotypic and genomic parallelism has been recently identified for many iconic examples of parallel evolution in animals, such as threespine sticklebacks (Stuart et al., 2017; Magalhaes et al., 2021), salmonid fishes (Jacobs et al., 2020), guppies (Whiting et al., 2021), cichlids (Weber et al., 2021), marine snails (Morales et al., 2019), and songbirds (Salmón et al., 2021), though only rarely investigated in plants (but see Knotek et al., 2020; James et al., 2021). For instance, Bohutínská et al. (2021) showed a divergence-dependent level of genomic parallelism in Brassicaceae; i.e., the degree of gene reuse was decreasing with the increasing divergence between lineages. This is in congruence with findings in *Senecio lautus* (James, Arenas-Castro, et al., 2021; James, Wilkinson, et al., 2021), in which similar phenotypes with large divergence times within-population pairs have arisen via mutational changes in different genes, although many of these genes shared the same biological functions. However, even in the system of a recent post-glacial origin, such are the alpine populations of *Heliosperma pusillum* (Trucchi et al., 2017), similar phenotypes were also achieved via selection on different genes in the same functional pathways. In such polygenic systems we expect genetic redundancy; i.e., mutations in different genes (possibly within the same pathway) can lead to similar phenotypes and thus to phenotypic parallelism (Hermisson and Pennings, 2017; Höllinger et al., 2019; Barghi et al., 2020). In systems with high heterogeneity among the compared ‘parallel’ habitat types such as alpine stands, which may strikingly differ in multiple local parameters (e.g., altitude, type of substrate, exposure), the extent of genomic and phenotypic parallelism may be underestimated. We thus need empirical evidence from other systems encompassing recent adaptation to narrowly defined environmental challenges to explore if genetic redundancy can lead to phenotypic parallelism.

Serpentines, naturally toxic edaphic substrates, represent one of the most challenging environments for plants. They are primarily distinguishable by markedly low soil concentrations of Ca, high concentrations of Mg (thus a highly skewed Ca/Mg ratio), and elevated heavy metals such as Cr, Co, and Ni (Brady et al., 2005; O’Dell and Rajakaruna, 2011). Consequently, unlike alpine or coastal environments, selective conditions of serpentine substrates can be easily replicated experimentally *ex situ*, as their specific soil chemistry is the principal defining factor. Although occurring worldwide, serpentine outcrops are typically scattered as small edaphic ‘islands’ surrounded by other types of substrates. Repeated colonization attempts from the surrounding non-extreme substrates then stimulate parallel evolution of serpentine adaptation, as has been recently shown in *Arabidopsis arenosa* (Konečná et al., 2021). In its genetically highly diverse autotetraploid populations, low neutral genetic differentiation and recent split times between geographically proximal serpentine and non-serpentine populations indicate recent postglacial serpentine invasions that happened at least five times independently (Konečná et al., 2021). Taken together with the described genetic basis of serpentine adaptation, *Arabidopsis arenosa* represents a suitable plant model for studying the repeatability of adaptation towards a well-defined selective environment. Yet, the extent of phenotypic parallelism across relevant fitness and physiological traits and the question of whether both levels correspond in the extent and magnitude of parallelism remained unknown.

Here, we used reciprocal transplants within three serpentine – non-serpentine *A. arenosa* population pairs and asked, to what extent the phenotypic evolution is repeatable during rapid parallel edaphic adaptation. We hypothesize that in this recently established system with pervasive sharing of adaptive polymorphism among populations (Konečná et al., 2021), genetic and phenotypic parallelism will be largely congruent. By coupling available population genomic data with transplant experiments in the same populations, we quantified (i) genomic parallelism at the level of genes (shared selection candidate loci) and functional pathways (shared gene ontology categories) and (ii) parallel phenotypic changes by scoring a diverse set of traits: fitness proxies, morphological functional traits, and ion uptake. We specifically asked: (i) Do serpentine populations exhibit fitness advantage in their local serpentine soil? If so, does the direction and magnitude of fitness response differ among population pairs in congruence with the level of their neutral genetic differentiation? (ii) Is there an evolutionary cost of serpentine adaptation (trade-off) and how does its extent vary over the three parallel population pairs? (iii) Does the extent of phenotypic parallelism correspond to gene-level parallelism, or is the gene parallelism limited due to genetic redundancy?

Local adaptation has been manifested in a congruent direction across all three parallel serpentine populations and resulted in the soil-of-origin advantage of serpentine populations and only very limited trade-offs. However, the magnitude of the adaptive response differed, reflecting the level of genetic differentiation within-population pairs. Phenotypic parallelism in functional traits was pervasive, only a minority of traits showed non-parallel variation. Contrary to our initial hypothesis, gene-level parallelism was rare, especially when compared to high similarity in functional pathways. As a consequence, similar phenotypes could arise mainly via selection on different genes in similar genetic pathways, highlighting the role of genetic redundancy and stochasticity in rapid adaptation with polygenic basis.

## MATERIAL AND METHODS

### Study species and environment

*Arabidopsis arenosa* is an obligate outcrosser that is widespread across most of Europe on semi-open rocky habitats characterised by reduced competition and grows on various substrates, predominantly on calcareous and siliceous soils (Schmickl et al., 2012). For this study we collected seeds from three autotetraploid serpentine populations (S1, S2, S3) and three geographically proximal (<19 km apart) sister autotetraploid non-serpentine siliceous populations in Central Europe (N1, N2, N3; STable 1; Konečná et al., 2021). The differences in elevation within the pairs were small for S1-N1 – 71 m, S2-N2 – 22 m, and greater for S3-N3 – 322 m, however, the variation in local vegetation structure (open mixed/coniferous forests with rocky outcrops) and in climatic conditions among all sites was negligible (Konečná et al., 2021). In contrast, we observed strong differences in soil chemistry, mainly driven by a high level of heavy metals (Co, Cr, and Ni), high concentrations of Mg, and low Ca/Mg ratios consistently differentiating serpentine substrates occupied by the studied populations from their paired adjacent non-serpentine counterparts (see Konečná et al., 2021 for details and SFig. 1). Thus, we consider substrate as a primary source of selective divergence between replicated serpentine and non-serpentine populations and use different soils as experimental treatments in the analysis of serpentine adaptation.

### Experimental cultivations

To test for local substrate adaptation, we compared the plant performance in native versus foreign soil using a reciprocal transplant experiment. We reciprocally transplanted plants of serpentine and non-serpentine origin within three pairs of adjacent populations (S1–N1, S2– N2, S3–N3) that served as representatives of independent serpentine colonisations (Konečná et al., 2021). For each population pair, we transplanted young seedlings into serpentine and non-serpentine soils sampled at the original sites (i.e., S1 plants were cultivated in both S1 and N1 soils), ∼24 plants per population and treatment, 284 plants in total. We cultivated them in the experimental garden of the Faculty of Science, Charles University with similar climatic conditions as in the original sites (methodological details are provided in Supplementary Methods) and tested for the substrate of origin*soil treatment in selected fitness indicators (described further) as a proxy of local substrate adaptation. We observed initial differences already in germination and young rosette sizes (Konečná et al., 2021), and thus continued with the cultivation for the entire growing season until seed set.

To analyse the early plant response to altered Ca/Mg ratios in serpentine vs. non-serpentine soils, we conducted additional *in vitro* experiment focused on quantifying root growth in seedlings. Similarly to Berglund et al. (2004) we focused on root phenotypes, which directly mirror the stressful solutions, and they are visible even in the early stages of plant development. We germinated and cultivated plants *in vitro* (five seedlings per Petri dish) on agar-solidified media supplemented with various Ca/Mg ratios – 1.97 in the control medium, 0.2 in medium reflecting the mean Ca/Mg ratio across our studied serpentine populations, and 0.04 in medium according to Bradshaw (2005). The experiment was conducted in three cultivation batches, altogether 549 seedlings were cultivated. For cultivation details see Supplementary Methods.

### Trait scoring

The extent of phenotypic parallelism may depend on the choice of measured traits and some of these, e.g., physiological or early developmental traits, are only rarely evaluated due to technical limitations. Here, we aimed to capture a broad range of phenotypic variation by scoring a diversity of potentially relevant traits. Specifically, on plants cultivated in soils we scored bolting and flowering time, rosette area, number of additive rosettes as morphological life-history traits, further, fruit production, total seed mass production, above-ground and root biomass and survival as fitness proxies, and ion uptake of five important elements characterizing serpentine substrate (Co, Cr, Ni, Ca, Mg) (SData 1 and SData 2). In seedlings grown on media, we scored additional functional traits of root growth and architecture: total root growth, main root length, and density of lateral roots (SData 3). This suite of phenotypic traits allowed us to cover both early and later stages of plant development, and macroscopic as well as physiological properties.

Plants were cultivated for 10 months (January – October 2019). Four months after germination, leaf rosettes of plants reached their maximum diameter (SFig 6 in Konečná et al., 2021) and we measured rosette areas. We scored the time of bolting when the stems started elongating (at least 1 cm tall) and the first flowering bud appeared, and flowering time as the time of opening of the first flower. Mature fruits were collected in regular intervals 1-2 times a week since mid-June and then also synchronously at two-time points when they were produced by most of the individuals. Fruits containing less than five seeds were considered not well developed and were excluded (mean number of seeds/fruit: 73). We provide a summary of flowering plants and fruit proportion of plants in STable 2. We counted and weighted seeds per plant, which were collected continuously (1-22 fruits/plant). We calculated average seed mass per fruit to calculate the total seed mass production by multiplying this value the total number of developed fruits, assessed by fully developed diaphragmas. To account for a possible seasonal variability in seed mass, we compared the seeds produced by the end of July with those produced later during summer and autumn, but no significant difference was observed (results not shown). In early September, we counted the total number of additive rosettes that were produced by the plants that reflect the potential of the plants to survive and further reproduce in the next year. The experiment was terminated at the end of October, in the end of main vegetation season, we harvested the plants and weighted dried above-ground biomass (all surviving plants) and root biomass (5 randomly chosen surviving plants/population/treatment due to challenging and time-consuming extraction). Plant survival was surveyed continuously until the end of the experiment. Finally, we measured several root traits (total root growth – length of all roots, main root length, and density of lateral roots) in plants 15 days since germination on agar media.

We measured uptake of the key serpentine-distinctive elements (Ca, Co, Cr, Mg, and Ni) into above-ground tissues in randomly selected 10 individuals from each study population when cultivated in serpentine-soil treatment. After 2.5 months of cultivation in serpentine soils, leaf samples were collected, rinsed with distilled water, air-dried and decomposed prior to their analysis (see Supplementary Methods for details). To identify potential contamination of leaf samples by heavy metals (Co, Cr, and Ni), we used the 1.5×IQR rule to detect outlier values (i.e., exceeding 1.5 × interquartile range of elemental concentrations) and the resulting four samples (6% of data) were removed from all analyses.

### Leaf and soil elemental profiles

Based on elemental composition of soil in our studied natural populations from Konečná et al. (2021), we selected five elements contributing the most to the serpentine soil-induced stress (i.e., Ca, Co, Cr, Mg, and Ni) and monitored their concentrations in the reciprocal transplant experiment, both in the substrate for cultivation and directly in plant leaf tissues. Soil samples were collected in mid-May from five pots/population/treatment, then mixed, sieved and dried at 60 °C. Leaf samples were taken at the same time. Both soil and leaf samples were processed according to a protocol summarised in Supplementary Methods. Concentrations of elements were quantified using inductively coupled plasma mass spectrometry (ICP-OES; spectrometer INTEGRA 6000, GBC, Dandenong Australia).

### Statistical analysis of trait variation

We assessed the variation in morphological life-history traits and fitness proxies by linear and generalized linear models (LM and GLM) and in functional root traits by linear mixed effect models (LMM). For each trait separately, we tested the (fixed) effects of native substrate of origin (S vs. N), population pair (1, 2, 3), treatment (S vs. N), and interaction substrate of origin*treatment. Traits deviating from normality were log-transformed or square-root transformed. In LMMs, we used random effects of cultivation batch and Petri dish. No pair of measured traits was highly correlated (R > 0.8, SFig. 2). Tests of significance for particular fixed-effect factors and interaction substrate of origin*treatment were conducted using Type III Wald χ2 tests in the R package car ver. 3.0-12 (Fox and Weisberg, 2019).

We used aster models (Geyer et al., 2007; Shaw et al., 2008) to analyze the hierarchical composite fitness differences among populations separately within each soil treatment. This approach is suitable for combining multiple fitness traits with different distributions into a single composite fitness variable. For each individual, we used the following hierarchy: 1 → survival and transition to bolting (Bernoulli distribution, 0/1) → survival and transition to flowering (Bernoulli distribution, 0/1) → survival and transition to fruit production (Bernoulli distribution, 0/1) → total seed mass production (Poisson distribution). By comparing the nested models with the effect of substrate of origin, population pair, and their interaction using likelihood ratio tests, we found the model with the highest likelihood (substrate of origin*population pair) (STable 3). Using likelihood ratio tests, we also tested the differences between populations within population pairs in order to assess if populations significantly differ in composite hierarchical fitness.

To further quantify fitness response using a single value of cumulative fitness taking into account all fitness proxies scored over the entire vegetation period, we multiplied flowering proportion and early survival, fruit production, total seed mass production, late survival after fruit production, and above-ground biomass for both serpentine and non-serpentine plants in their native and foreign soil. The values of total seed mass production and above-ground biomass were standardized by the average over all plants (regardless of substrate origin) grown in the control non-serpentine treatment for each pair separately to account for variation between distinct genetic lineages, i.e. population-pairs. Besides the phenological and reproductive traits, we also included late survival and above-ground biomass, proxies for the plant’s success in the following year. Then, we used cumulative fitness to quantify the overall fitness trade-offs (cost of adaptation; following Hereford, 2009). We calculated the difference in relative fitness between a pair of serpentine and non-serpentine populations in each population’s native soil. These estimates were standardized by relative mean fitness at each soil. Further, we evaluated the relative magnitudes of local adaptations to serpentine soils by comparing serpentine and non-serpentine plants within each population pair cultivated in serpentine soil (following Hereford, 2009). The magnitudes of adaptations were expressed in terms of differences between the substrate of origins in each population pair. We calculated these values based on overall cumulative fitness estimates. The difference (native – foreign) represents the total increase in fitness due to local adaptation (if the values are positive).

We tested the differences in morphological life-history traits, fitness proxies, and leaf elemental concentrations (ionomic phenotypes) between the group of serpentine and non-serpentine populations cultivated in the stress environment (serpentine soil) using LM with substrate of origin, population pair and their interaction as fixed effects. To assess the degree of parallelism and non-parallelism more quantitatively, effect sizes of each factor and their interaction were estimated from the output of LM models (trait ∼substrate of origin + population pair + substrate of origin*population pair) using a partial eta-squared method (anova_stats function) in the R package package sjstats ver. 0.18.1 (Lüdecke, 2021). Larger effect of the substrate of origin than the effect of the substrate of origin*pop. pair interaction was an indication of parallelism while the opposite implied non-parallel serpentine/non-serpentine differentiation in that particular trait (Stuart et al., 2017).

### Quantification of genomic parallelism

To quantify gene-level parallelism, i.e. the proportion of shared candidate loci out of all differentiation candidate genes, we used the already published individual resequencing data for the serpentine (S1, S2, and S3) and non-serpentine (N1, N2, and N3) populations (7-8 individuals per population, 47 in total) from Konečná et al. (2021). Raw data are available as BioProject PRJNA667586. To quantify the extent of gene parallelism we leveraged the threefold-replicated natural setup and already identified candidate genes that show repeated footprints of selection across the three events of serpentine colonization in Konečná et al. (2021). Briefly, differentiation candidates were identified as gene-coding loci exhibiting excessive differentiation between serpentine – non-serpentine populations within each population pair using 1 % outlier F_ST_ window-based scans (SData 4). Further, we identified parallel differentiation candidates as overlapping genes in differentiation candidate lists across at least two population pairs. We tested if such overlap is higher than a random number of overlapping items given the sample size using Fisher’s exact test in SuperExactTest in R package ver. 1.0.7 (Wang et al., 2015). Further, we calculated a proportion of shared genes as number of shared differentiation candidate genes per each two population pairs divided by the number of remaining ‘unique’ (non-shared) differentiation candidate genes from both populations from that pair.

To infer function-level parallelism, we annotated differentiation candidates into biological processes, molecular pathways, and cellular components using gene ontology (GO) enrichment analysis for each population pair separately (SData 5). We applied Fisher’s exact test using the ‘classic’ algorithm implemented in R package topGO ver. 2.38.1 (Alexa and Rahnenfuhrer, 2021). We worked with *A. thaliana* orthologs of *A. lyrata* genes obtained using biomaRt and *A. thaliana* was also used as the background gene universe in all gene set enrichment analyses. With the ‘classic’ algorithm each GO term is tested independently and it is not taking the GO hierarchy into account. Therefore, it is more suitable for comparisons across multiple lists of candidates to get the overall quantification of parallelism. We retained only significant GO terms (p < 0.05, Fisher’s exact test). We calculated the proportion of shared functions as the number of shared GO terms per each two population pairs divided by the number of remaining ‘unique’ (non-shared) GO terms for selected population pair combination. In order to verify that functional parallelism in GO terms is not driven solely by the parallel differentiation candidate genes, we ran the same analyses also without these candidates.

Finally, we searched for functional associations among the identified differentiation candidates from three population pairs using STRING v11 database (Szklarczyk et al., 2015) of protein-protein association networks. We used ‘multiple proteins’ search in *Arabidopsis thaliana* and these active interaction sources: co-expression, co-occurrence, databases, experiments, gene fusion, neighborhood, and textmining. We required the minimum interaction score of medium confidence (0.4) and retained only 1st shell associations.

## RESULTS

### Varying magnitude and direction of parallel serpentine adaptation in Arabidopsis arenosa

In line with the hypothesis of parallel adaptation, originally serpentine populations grown in their native soils exhibited higher values than non-serpentine plants in most of the fitness-related functional traits. Significant interaction of the substrate of origin*treatment (*p* < 0.05, Type III Wald χ2 tests, Table 1, SFig. 3) was observed for the rosette area, number of additive rosettes, total seed mass production, above-ground and root biomass, and all three root traits. To further quantify the overall fitness effects, we modelled the joint contribution of four reproductive plant traits in hierarchical aster models, which explicitly account for the inter-dependence between particular reproductive fitness traits and their transitions within each treatment (Fig. 1A). We observed significantly higher composite fitness values for originally serpentine populations S1 and S3 grown in their native serpentine substrate as compared to their paired non-serpentine populations (Fig. 1A; 45.63 and 56.42, respectively; STable 4). The difference in population pair 2 was negligible, reflecting the overall very low survival of both populations (62 %) in S2 serpentine soil (STable 2).

**Table 1.** Significance of the effects of substrate of origin (S, serpentine vs. N, non-serpentine), population pair (1, 2, 3), treatment (S vs. N soil) and the substrate of origin*treatment interaction on functional traits of *Arabidopsis arenosa* individuals cultivated in a reciprocal transplant setting. Differences were evaluated using linear models (LM), binomial generalized linear models (GLM binomial), and linear mixed effect models (LMM). Tests of significance for individual fixed effect factors and interaction were conducted by Type III Wald χ2 tests. The random effects in LMM were cultivation batch and Petri dish for cultivation. The traits have been measured in a reciprocal transplant experiment with manipulated soil, except for the last three traits that were scored in an additional *in vitro* experiment using growth media with manipulated Ca/Mg concentrations).

| Response variable | Trans formation | Model | Substrate of origin |  |  | Pop. pair |  |  | Treatment |  |  | Substrate of origin*treatment |  |  |
| --- | --- | --- | --- | --- | --- | --- | --- | --- | --- | --- | --- | --- | --- | --- |
| | | | Df | $\chi^2$ | *p* | Df | $\chi^2$ | *p* | Df | $\chi^2$ | *p* | Df | $\chi^2$ | *p* |
| Bolting time [days] | log | LM | 1 | 2.0524 | n | 2 | 26.3506 | *** | 1 | 6.8092 | ** | 1 | 2.4737 | ns |
| Flowering time [days] | log | LM | 1 | 0.0088 | ns | 2 | 3.5704 | * | 1 | 5.5588 | * | 1 | 0.0618 | ns |
| Rosette area [mm<sup>2</sup>] | sqrt | LM | 1 | 74.0269 | *** | 2 | 4.0045 | * | 1 | 382.8176 | *** | 1 | 51.43337 | *** |
| N of additive rosettes | log | LM | 1 | 34.4439 | *** | 2 | 9.9119 | *** | 1 | 91.4885 | *** | 1 | 14.2435 | *** |
| Probability of fruit production (0/1) |  | GLM binomial | 1 | 10.2295 | ** | 2 | 3.576 | ns | 1 | 14.0429 | *** | 1 | 2.8821 | . |
| Total seed mass production [mg] | log | LM | 1 | 27.2423 | *** | 2 | 6.3113 | ** | 1 | 26.9319 | *** | 1 | 12.6945 | *** |
| Above-ground biomass [mg] | sqrt | LM | 1 | 9.6231 | ** | 2 | 9.6957 | *** | 1 | 104.7775 | *** | 1 | 13.8185 | *** |
| Root biomass [mg] | log | LM | 1 | 15.6522 | *** | 2 | 16.9575 | *** | 1 | 61.3215 | *** | 1 | 14.2662 | *** |
| Probability of late survival (after fruit production) (0/1) |  | GLM binomial | 1 | 6.5438 | * | 2 | 4.7977 | . | 1 | 8.0743 | ** | 1 | 2.1933 | ns |
| Total root growth [cm] | log | LMM | 1 | 123.7103 | *** | 2 | 14.7846 | *** | 2 | 60.0113 | *** | 2 | 44.4136 | *** |
| Main root length [cm] |  | LMM | 1 | 16.4607 | *** | 2 | 8.9317 | * | 2 | 24.9806 | *** | 2 | 6.2092 | * |
| Density of lateral roots (number of lateral roots/main root length) | sqrt | LMM | 1 | 12.9772 | *** | 2 | 17.5417 | *** | 2 | 20.2030 | *** | 2 | 8.0637 | * |

**Fig. 1.**
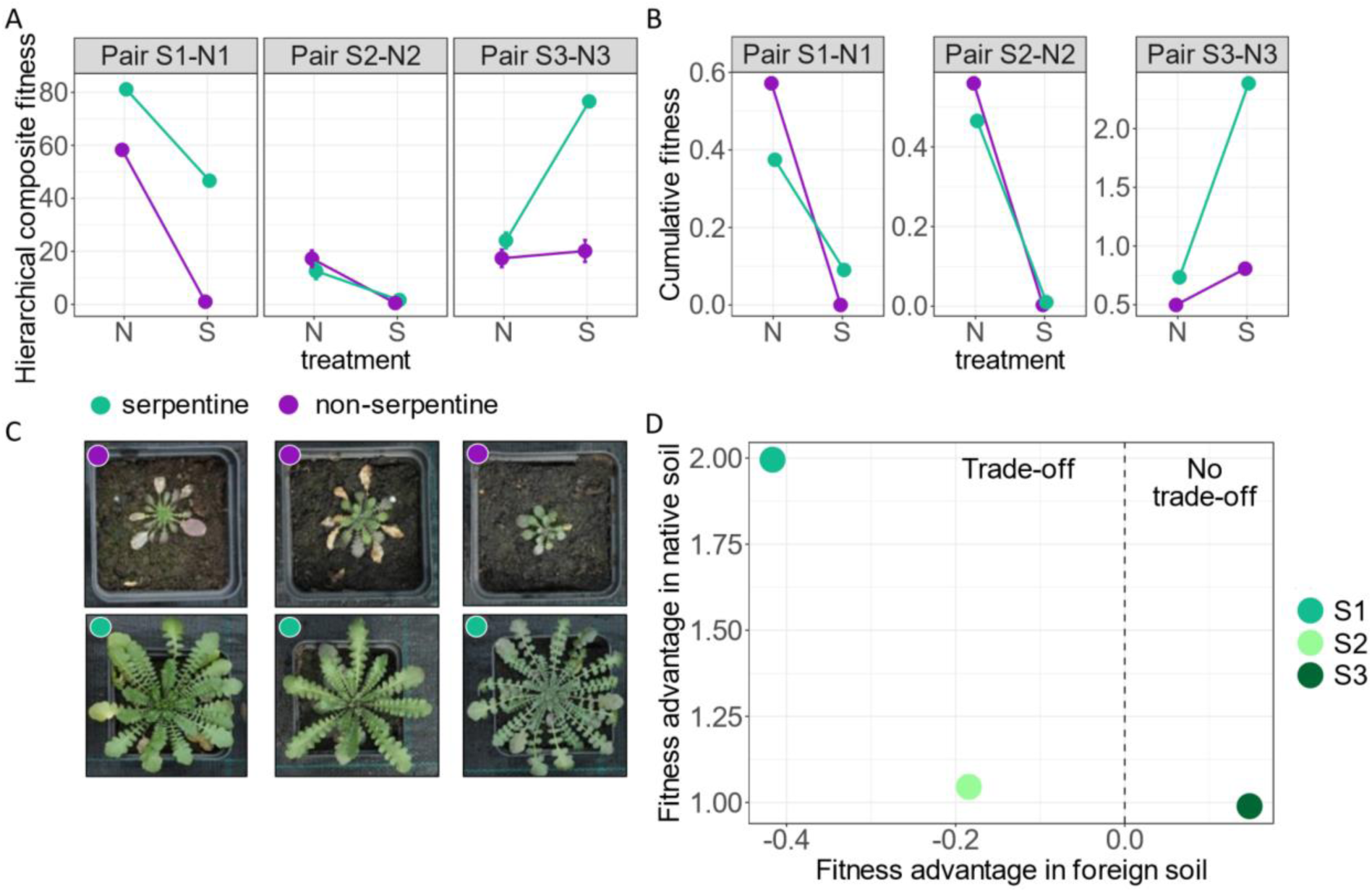
Local adaptation to serpentine soils in three parallel population pairs of *A. arenosa*. A) Hierarchical composite fitness (+-SEs) for serpentine (green) and non-serpentine (violet) plants cultivated in local serpentine (S) and non-serpentine (N) soils. Estimates were taken from hierarchical aster models combining proportion of bolting plants, proportion of flowering plants, fruit production, and total seed mass. The significant (p < 0.001) interactions were found for the S1 - N1 and S3 - N3 population pairs (complete results in STable 4). B) Cumulative fitness was calculated by combining five reproductive and vegetative traits (flower proportion and early survival, fruit production, total seed mass production, late survival after fruit production, and above-ground biomass). Note the y-axes have been adjusted to 4 months of cultivation (dot colours denote the plant origin; from the left: population pair 1 – 2 – 3; photos taken by V. Konečná). D) Relative fitness advantage of originally serpentine populations in their native and foreign (i.e. non-serpentine) soils calculated based on cumulative fitness estimates. On the left side, both populations from the pair (due to our focus on serpentine adaptation only serpentine populations are visualized) had higher relative fitness in their native soils indicating fitness trade-off. On the right side, the serpentine population S3 had relatively higher fitness than non-serpentine population in both soil treatments. Note: each fitness advantage estimate was standardized by the relative mean fitness of plants at each soil treatment.

Further, we calculated cumulative fitness (STable 5) based on vegetative and reproductive fitness proxies, to quantify differences in the magnitude of local adaptation among population pairs (Fig. 1B). In all cases, the magnitude of local adaptation was positive (Table 2), which indicates the advantage of serpentine populations in their native soil and selection against immigrants (Hereford, 2009). Yet, the magnitude of this response differs among the three population pairs, which is in line with varying relative fitness differences observed in aster models. We found the lowest value for population pair 2 and highest for pair 3, which is fully in congruence with increasing genome-wide differentiation of population pairs 2 < 1 < 3 (Table 2). Overall, serpentine populations have higher fitness in serpentine soils than their originally non-serpentine counterparts, although its magnitude strongly varied, altogether providing strong evidence for substrate-driven local adaptation.

**Table 2.** Estimated magnitude of local adaptation in serpentine soils.

| Population pair | Genetic differentiation ($F_{ST}$)<sup>1</sup> | Substrate environmental distance<sup>2</sup> | Magnitude of local adaptation — cumulative fitness<sup>3</sup> | Relative difference in hierarchical composite fitness<sup>4</sup> |
| --- | --- | --- | --- | --- |
| S1-N1 | 0.069 | 4.174 | 0.090 | 45.635 |
| S2-N2 | 0.029 | 6.682 | 0.007 | 1.129 |
| S3-N3 | 0.085 | 6.934 | 1.579 | 56.414 |

<sup>1</sup> inferred from genome-wide nearly-neutral fourfold degenerated single nucleotide polymorphisms (SNPs) by Konečná et al. (2021)
<sup>2</sup> substrate environmental distances were calculated based on chemical analysis of 20 elements in soils sampled at original locations
<sup>3</sup> cumulative fitness gathered by multiplication of flower production and early survival, fruit production, total seed mass production, late survival after fruit production, and above-ground biomass
<sup>4</sup> inferred by hierarchical aster model performed on proportion of bolting plants -> proportion of flowering plants -> fruit production -> total seed mass production

In line with differences in the direction of the substrate of origin*treatment interaction we also found variation in trade-offs. While S1 and S2 populations exhibited costs of serpentine adaptation in the non-serpentine soils (based on cumulative fitness values), fitness advantage in both soil types was present in the S3 population (0.148) (Fig. 1D). Yet, even in the cases involving trade-offs, serpentine adaptation had a limited cost (1.52 average fitness advantage in native soil vs. -0.3 average fitness advantage in foreign soil, Fig. 1D). This is also in congruence with only subtle and nonsignificant differences in the survival rate of plants in non-serpentine treatment (89 % and 87 % for plants of serpentine and non-serpentine origin, respectively).

### Varying levels of phenotypic parallelism across functional traits

We further analyzed the magnitude of response to serpentine for particular functional traits across distinct population pairs in order to identify traits exhibiting parallel responses. We identified traits exhibiting strong parallel response to the serpentine soil as those having a larger effect of the substrate of origin than of the substrate of origin*population pair interaction when grown in the selective serpentine soil (Fig. 2A, B). Regardless of the population pair, originally serpentine plants bolted earlier, had larger rosette area, higher number of additive rosettes, higher seed mass, higher root biomass, and lower uptake of Ca, Co, Cr, Mg, and Ni than non-serpentine plants when cultivated in serpentine soil (STable 6, Fig. 2A, B, SFig. 4).

**Fig. 2.**
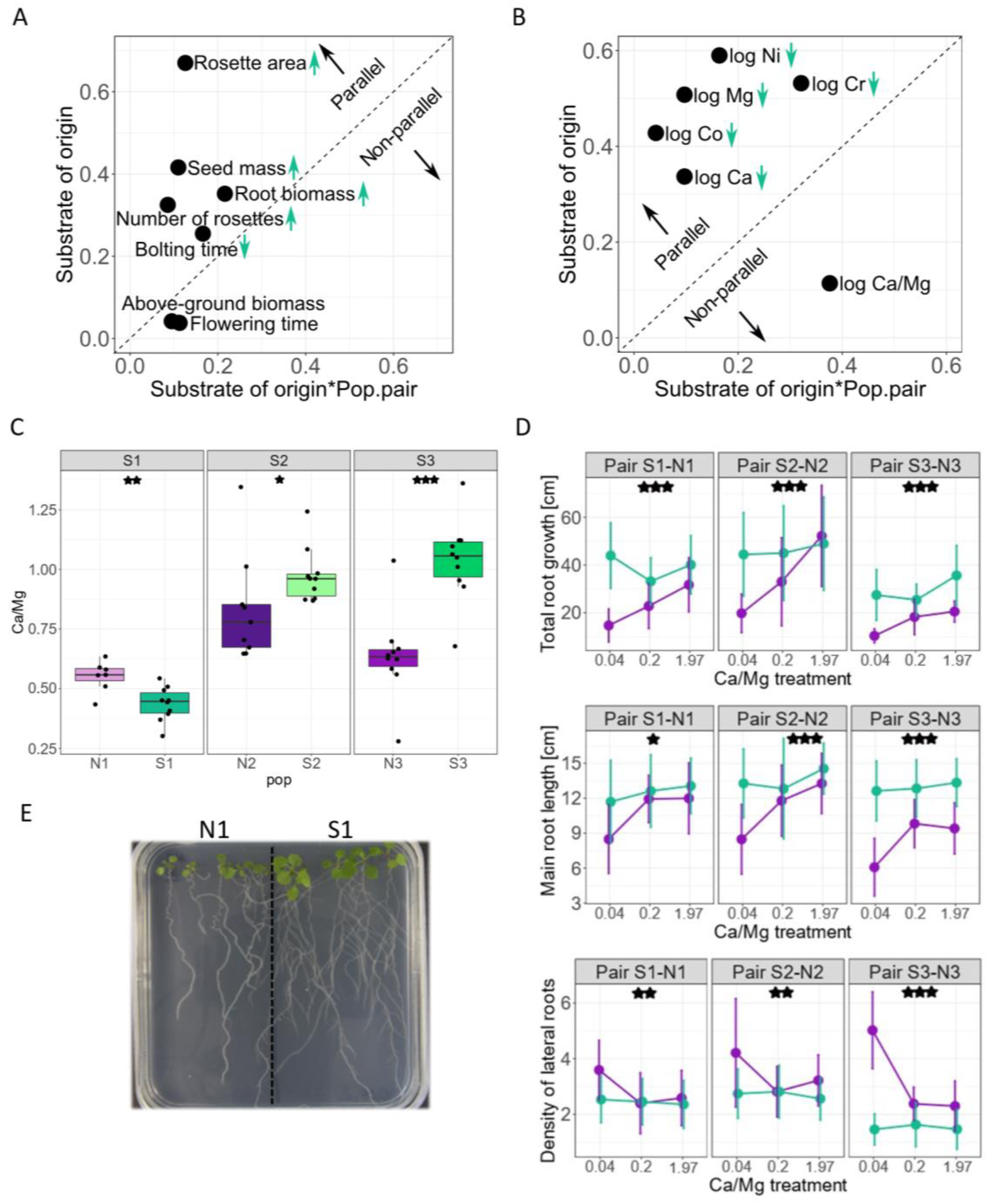
Variation in the extent of parallelism among particular morphological life-history traits (A) and leaf elemental concentrations (B) scored on plants cultivated in the selective serpentine soils. The extent of parallelism was estimated as effect sizes (Eta-squared) in linear models addressing the effect of the substrate of origin (S, serpentine vs. N, non-serpentine), pop. pair (1, 2, 3), and their interaction, calculated separately for each trait (see STable 7 for input values). The effect size of the substrate of origin (y-axis) shows the extent to which a given trait diverges predictably between serpentine and non-serpentine plants, i.e., in parallel, while the substrate of origin*population pair effect size (x-axis) quantifies the extent to which serpentine/non-serpentine phenotypic divergence varies across population pairs (i.e., deviates from parallel). Points falling above the dashed line have a larger substrate of origin effect (i.e. parallel) than the substrate of origin*pop. pair interaction effect (i.e. non-parallel). Green arrows indicate the overall trend in trait value in populations of serpentine origin. C) Variation in the ratio of Ca/Mg content in the leaves of paired serpentine and non-serpentine populations cultivated in their corresponding serpentine soils. D) Differences in root growth of the three population pairs after 15 days since germination; note: 1.97 is a control medium, green: plants of serpentine origin, violet: plants of non-serpentine origin. E) Illustrative photo from the cultivation of N1 and S1 plant in 0.04 Ca/Mg treatment. Asterisks denote the significance of the effect of substrate of origin*treatment interaction (*** *p* < 0.001; ** *p* < 0.01; * *p* <= 0.05; STable6).

On the contrary, we found only a minority of traits that showed rather a non-parallel variation, i.e., a dominant effect of the substrate of origin*population pair interaction: above-ground biomass, flowering time, and Ca/Mg ratio in plants. For flowering time, only in population pair 3, serpentine population flowered earlier than the non-serpentine one in the serpentine treatment (SFig. 3). The most remarkable difference was found for the Ca/Mg ratio, where S2 and S3 populations had relatively higher Ca/Mg ratios than their paired non-serpentine populations, while we observed a reverse trend in population pair 1 with lower (i.e. more skewed) values in S1 (Fig. 2C; as a consequence of the lower Ca uptake, Fig. S2). To test if such differences in Ca/Mg uptake also correspond to non-parallel response specifically to altered Ca/Mg ratio in the environment, we cultivated plants of all populations *in vitro* on agar-solidified media with varying Ca/Mg ratios and scored root growth and architecture. We observed consistently better root growth of originally serpentine over non-serpentine plants in highly skewed Ca/Mg environments (Fig. 2D, E). In addition, lower Ca/Mg ratio affected the root architecture: serpentine plants had longer main roots and a lower density of lateral roots. Interestingly, such results were consistent over the three population pairs (Fig. 2D), suggesting that the specific pattern of Ca and Mg uptake of S1 population in serpentine soil is not mirrored in distinct fitness response to skewed Ca/Mg concentrations *per se* and this population likely evolved a distinct mechanism how to cope with low Ca/Mg ratio in its above-ground tissues.

Altogether, in line with our initial expectations, the recently diverged serpentine populations exhibited pervasive phenotypic parallelism in functional traits apparent in both early and later developmental stages, and only a minority of traits showed non-parallel variation.

### Genomic drivers of phenotypic parallelism

To uncover the genetic architecture underlying the observed parallel phenotypes, we also quantified and characterized parallelism at the level of genes and functional pathways. First, we took differentiation candidate genes (1 % outliers in F_ST_ genetic differentiation between proximal pairs of S and N populations; SData 4) identified by Konečná et al. (2021) and tested for gene-level parallelism. Although we found a significant over-representation of shared genetic candidates across all combinations of population pairs (Fig. 3A; Fisher’s exact test; *p* < 0.05; SData 4), the proportion of shared candidate genes was rather low, ranging from 2.21 % to 2.46 % (Table S8).

**Fig. 3.**
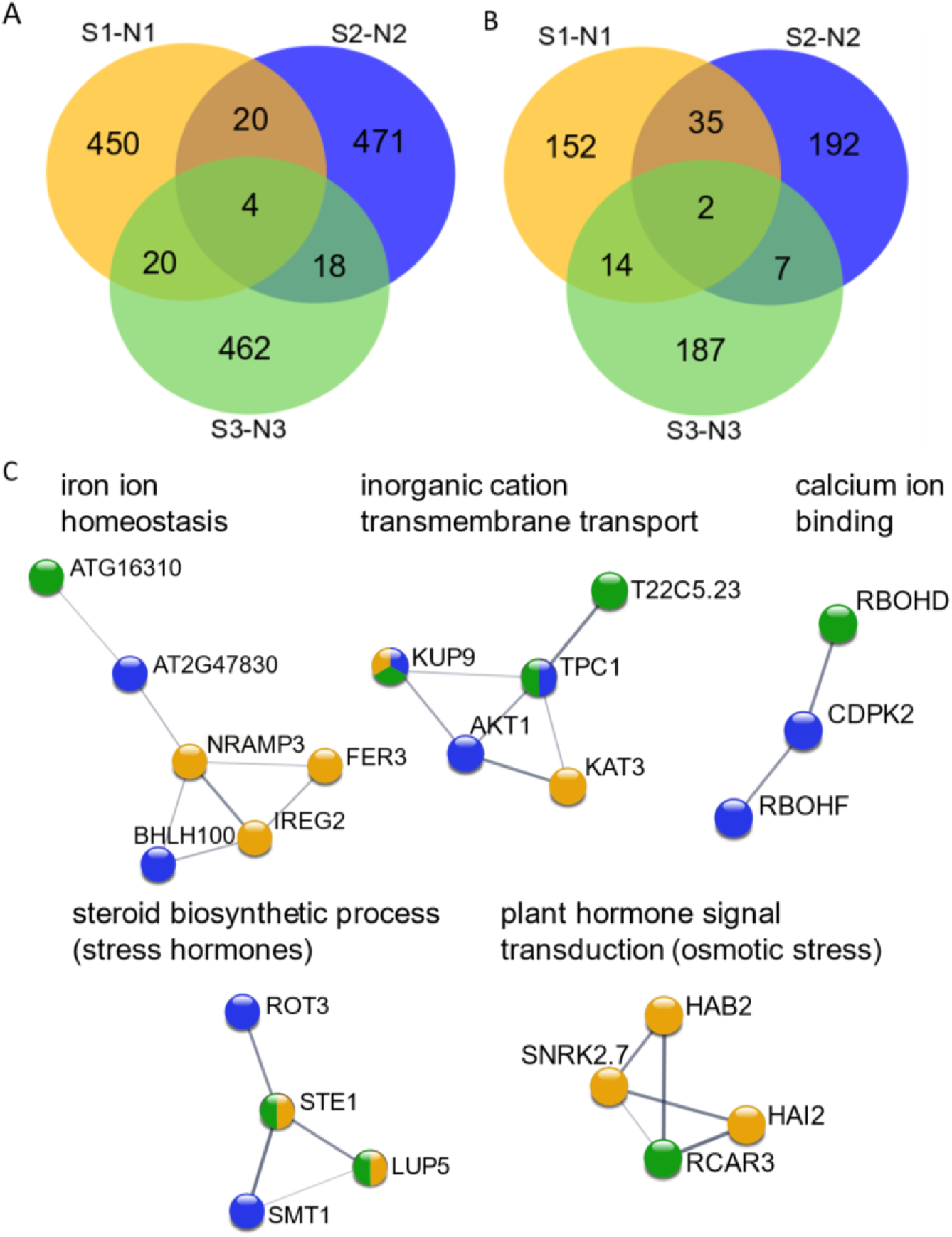
Significant genomic parallelism among the three investigated population pairs of *A. arenosa*. A) Venn diagram shows differentiation candidate genes identified in each population pair (1, 2, 3) and their intersection, i.e., parallel gene candidates. B) Venn diagram shows gene ontology terms in biological processes domain identified for each candidate gene list and their intersection over population pairs, i.e. parallelism by function. The number of overlapping items was significantly higher than random across all the intersections (*p* < 0.05; Fisher’s exact test; SData 4 and SData 5). C) Parallelism by function – significantly enriched biological processes or KEGG pathways extracted from STRING database; we used medium and higher confidence associations – displayed by the thickness of the lines (thin line = 0.4; thick line = 0.7). Genes identified as differentiation candidates in population pairs 1, 2 and 3 are marked by orange, blue and green colours, respectively. Parallel differentiation candidates are marked with more than one colour.

As the low gene-level parallelism might reflect genetic redundancy, we further associated all differentiation candidates identified for particular population pair with corresponding functional pathways (gene ontology – GO – enrichment analysis) and quantified function-level parallelism (Fig. 3B) as the percentage of shared enriched GO terms between any two pairs of populations (STable 8). All overlaps were significant (Fisher’s exact test; *p* < 0.05, SData 5) and the extent of pairwise overlap ranged from 2.06 % to 9.20 % for the most relevant category of biological process (BP; similar results were also achieved for molecular functions, and cellular components; STable 8). This relationship was not driven by the parallel differentiation candidate genes as we observed significant functional parallelism also when these genes were not used for GO enrichment (STable 8). Significantly enriched BP in all three population pairs were cellular process and metabolic process. BP categories shared between two population pairs were more specific, reflecting multiple processes that are relevant for facing the toxic serpentine soils (Fig. 4): (i) cellular transport: transport (shared between population pairs 2 and 3), protein transport, protein localization, and organonitrogen compound metabolic process (all three shared between pairs 1 and 2); (ii) abiotic stress: regulation of abscisic acid-activated signalling pathway, important in stress signalling (1 and 3), response to endoplasmic reticulum stress (1 and 2), and cellular response to stimulus (1 and 3); and (iii) developmental processes: anatomical structure development (1 and 2) and sexual sporulation (1 and 2) (for complete lists see SData 5).

**Fig. 4.**
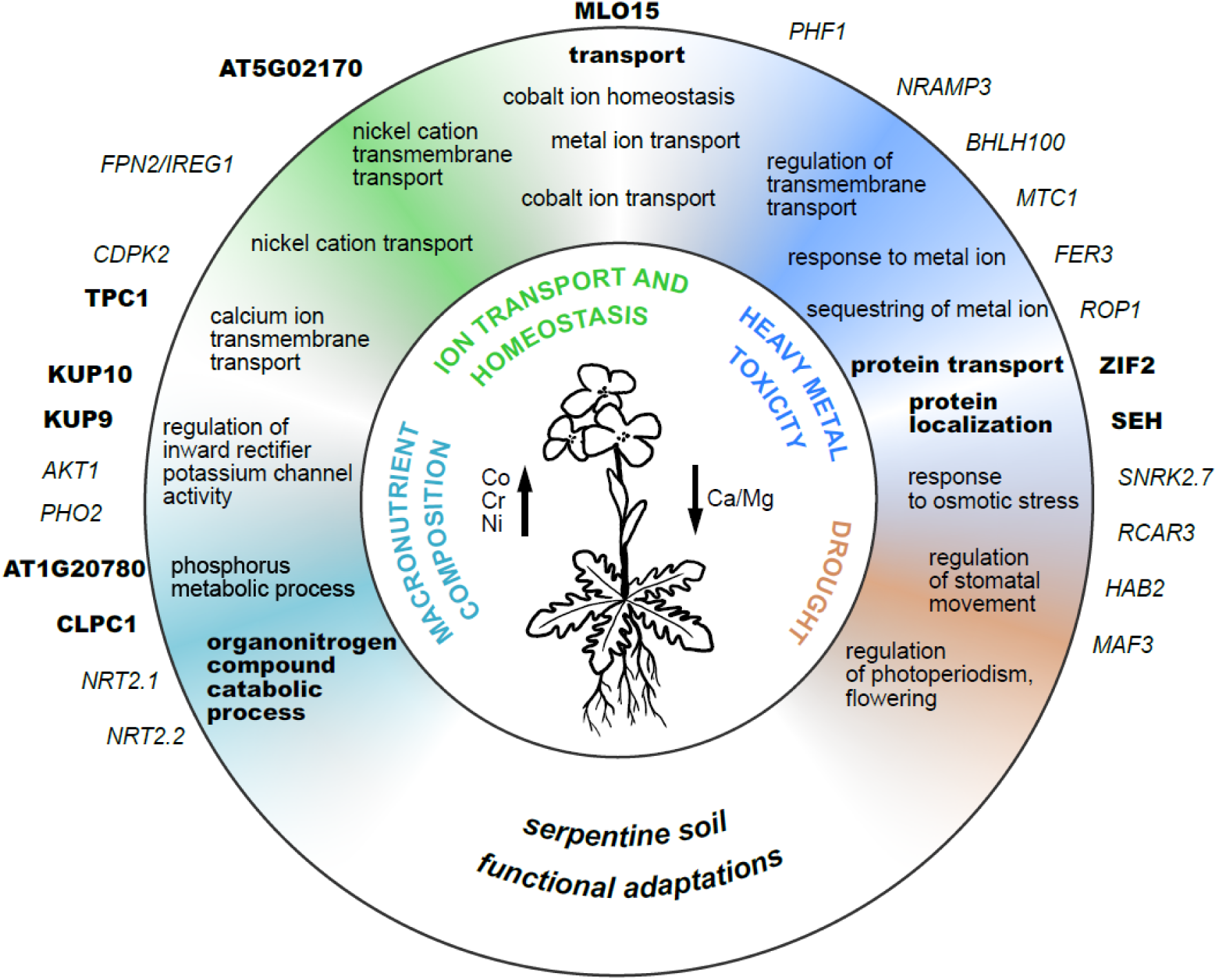
Functional responses to the major environmental challenges of serpentine substrates detected by gene and functional pathway-level analysis of *A. arenosa*. The figure summarizes key serpentine-specific challenges following (Brady et al., 2005; O’Dell and Rajakaruna, 2011; Konečná et al., 2020), relevant significantly enriched biological process GO terms and differentiation candidate genes which were inferred for at least one (thin) and two/three (i.e. parallel, in bold) serpentine populations.

To further inspect the function-level parallelism by specific pathway-oriented analysis, we inferred protein-protein functional networks from a combination of the differentiation candidates inferred for all three serpentine populations in STRING database. We found representative functional associations of distinct differentiation candidate genes inferred from different population pairs (Fig. 3C). For multiple functionally relevant parallel GO terms we showed that in each serpentine population, selection likely targeted different genes which belonged to the same pathway or developmental process. The STRING analysis also showed that candidate genes were highly inter-connected within such pathways (STable 9). Overall, we found significant parallelism by genes and functions associated with serpentine stress with evidence for selection on different genes from the same functional pathway.

## DISCUSSION

### Varying magnitude of local adaptation and limited trade-offs

Parallel fitness response in the direction of a home-site advantage detected in all three serpentine populations document overall significant local adaptation of *Arabidopsis arenosa* to toxic serpentine soils. However, the magnitude of fitness differences between originally serpentine and non-serpentine populations strikingly varied among the three population pairs, raising a question on the reason for such variation. Firstly, overall differences among particular selective environments may determine the strength of selection and thus also the extent of phenotypic divergence. A positive correlation between the magnitude of local adaptation and environmental distances, although with only a small effect has been frequently observed in plants (Hereford, 2009). Although the S3-N3 population pair exhibited the largest magnitude of local adaptation and was also the most diverged in substrate environmental distance, this was not true for the population pair S2-N2 with a comparable substrate environmental distance, yet by far the smallest magnitude of local adaptation response (Table 2). Secondly, neutral genetic differentiation, mirroring the extent of genetic divergence and intensity of gene flow, may indicate that population genomic processes play an important role (Savolainen et al., 2013). Indeed, the magnitude of local adaptation was in congruence with the genome-wide differentiation within population pairs in *A. arenosa*: higher genetic differentiation between geographically proximal serpentine and non-serpentine populations corresponded with a higher difference in fitness of plants cultivated in serpentine soils (Fig. 1A, B). Although there are multiple examples documenting local serpentine adaptation (reviewed e.g. in Brady et al., 2005; Konečná et al., 2020), experimental cultivations have not been accompanied by genetic investigations of population history leaving the generality of our finding unknown. As our system encompassed only three population pairs, studies on a larger number of S-N population contrasts are needed to systematically test the relationship between genetic differentiation and magnitude of local adaptation.

Using various combinations of fitness proxies, we detected utmost limited trade-offs of serpentine adaptation, i.e., minimal cost of serpentine adaptation in non-serpentine soil. Limited costs of adaptation seem to be in congruence with a major trend in other studies of local adaptation in general summarized by Hereford (2009). Generally, adaptation trade-offs can rather evolve in homogeneous environments (e.g., copper tolerance in *Mimulus guttatus* (Wright et al., 2013) than in heterogeneous environments (Bono et al., 2017)). For example, there was no difference in survival at non-serpentine sites between serpentine and non-serpentine *M. guttatus* (Selby and Willis, 2018) and only a little cost of adaptation was identified in serpentine *Arabidopsis lyrata* (Veatch-Blohm et al., 2017). Even in the pioneering serpentine reciprocal transplant experiments, Kruckeberg (1951) demonstrated the absence of substrate-driven trade-offs over various serpentine plants. He also proposed that competition can be the main cause of trade-offs. Although, there is generally not much evidence to support this hypothesis (Sianta and Kay, 2019), in *Helianthus exilis* local serpentine adaptation was observed only without competition (Sambatti and Rice, 2006). Due to the absence of competitors in our experiments, we cannot rule out that competition might add some cost of serpentine adaptation. On the other hand, similar habitat types, generally low-competitive open mixed/coniferous forests with rocky outcrops occupied by both serpentine and non-serpentine *A. arenosa* populations rather suggest comparable levels of competition in both habitats.

### Pervasive phenotypic parallelism

We detected parallel response to the selective serpentine substrate over the three independent serpentine populations in nearly all scored phenotypic traits (Fig. 2A, B). According to our expectations, in this recently diverged system (Konečná et al., 2021), parallelism is largely manifested at the phenotypic level. Traits with predominantly parallel responses accounted for 10 out of 13 all scored morphological life-history traits, fitness proxies, and traits related to ion uptake. Therefore, we show that the phenotypic trait divergence between serpentine and non-serpentine populations is largely predictable in *A. arenosa*. Experimental studies of serpentine adaptation revealed a range of life-history traits and physiological mechanisms showing similar responses among serpentine populations (e.g. Rajakaruna et al., 2003; Kolář et al., 2014), but also the variation in parallel traits depending on the specific chemistry of the serpentine soils being compared (e.g., Berglund et al., 2004). The overall extent of parallelism was rarely studied and comprehensively quantified across multiple morphological and life-history traits, and such studies focused mainly on animals (but see Knotek et al., 2020; James, Wilkinson, et al., 2021).

As specific soil chemistry is the principal selective factor on serpentines, regulation of micro-/macronutrients uptake and exclusion of heavy metals are expected to be the prime adaptive mechanisms (Brady et al., 2005; Kazakou et al., 2008). In line with this, serpentine substrate adaptation in *A. arenosa* is likely driven by high substrate content of heavy metals (Co, Cr, and Ni) and Mg, because these properties contributed the most to differentiating the original serpentine and non-serpentine soils (SFig. 1). Congruently, we found reduced uptake of all these elements in serpentine compared to non-serpentine plants when cultivated in serpentine soils, which was once again consistent across the three population pairs. *Arabidopsis arenosa* seems to reduce the uptake of heavy metals (such as Ni) as has been demonstrated for closely related *A. lyrata* (Veatch-Blohm et al., 2017) and some other plant species from the same location, such as *Knautia arvensis* group (Kolář et al., 2014). Importantly, plant response to the altered Ca/Mg ratios, perhaps the most distinctive character of serpentines (Brady et al., 2005), was apparent already in the early life stages. We observed a highly parallel trend of higher root growth of serpentine populations in media with highly skewed Ca/Mg ratios, well supporting similar studies on serpentine ecotypes of some other species (*Cerastium alpinum* and *K. arvensis;* Berglund et al., 2004; Kolář et al., 2014) yet not universally (e.g. *Galium valdepilosum;* Kolář et al., 2013 and *Streptanthus polygaloides;* Boyd et al., 2009). Similarly to *C. alpinum*, non-serpentine plants of *A. arenosa* had generally a higher number of lateral roots than serpentine plants. Different root growth strategies possibly evolved in serpentine populations as a response to high chemical stress, and the majority of resources are invested in the growth of the main root, while the formation of lateral roots is down-regulated (Berglund et al., 2004). Furthermore, the highest difference in this trait was in population pair 3, in which relevant differentiation candidate genes have been identified by a genomic divergence scan (reduce lateral root formation (*RLF*), repressor of lateral root initiation (*NRT2*.*1*)), suggesting a genetic basis for this phenotype.

### Causes and consequences of non-parallel phenotypes

In spite of generally pervasive phenotypic parallelism, we observed regionally specific patterns in three traits: above-ground biomass, flowering time and varying Ca/Mg ratio in leaves. On top of that, non-parallelism was also observed in the germination rate in serpentine soils in our previous study while N2 and N3 populations had 93 % and 86 % germination success on serpentine, respectively, seeds from N1 population did not germinate at all in serpentine soil (Konečná et al., 2021). Such locally-specific responses may reflect a complex interplay among distinct genetic basis of adaptive traits, genetic drift, and local environmental heterogeneity (Rosenblum et al., 2014; Fraïsse and Welch, 2019; James, Wilkinson, et al., 2021). In the case of the leaf uptake of Ca and Mg ratio, S2 and S3 populations accumulated relatively higher leaf Ca/Mg ratios than proximal non-serpentine populations, yet the opposite was observed for the S1-N1 population pair. The highly skewed Ca/Mg ratio (<1) is a major factor defining the serpentine soil stress worldwide (O’Dell and Rajakaruna, 2011) and the importance of Ca availability under increased Mg content has been highlighted for almost a century (Novák, 1928; Vlamis and Jenny, 1948; Vlamis, 1949; Kruckeberg, 1954; Walker et al., 1955). For instance, adapted serpentine populations of *A. millefolium* had higher leaf Ca/Mg ratios than non-serpentine plants, suggesting an efficient root to shoot transfer of calcium, selective uptake of Ca, or exclusion of Mg (O’Dell and Claassen, 2006). Variation in leaf Ca/Mg ratio among serpentine *A. lyrata* populations has been also shown by Veatch-Blohm et al. (2013). In *A. arenosa*, reduced uptake of Mg seems to be an important mechanism for regulating the Ca/Mg ratio in serpentine-adapted plants from populations S2 and S3 but not S1 (SFig. 4). Cellular concentrations of Ca^2+^ cations are controlled by an array of channels and carriers pointing to the complex genetic basis of Ca homeostasis and signalling (Tang and Luan, 2017). Interestingly, one of the parallel differentiation candidate genes encoding central calcium channel *TPC1* (Choi et al., 2014) appeared as a selection candidate in population pairs 2 and 3, but not in pair 1 (Konečná et al., 2021), suggesting that the observed non-parallelism may have a genetic basis.

The other non-parallel trait was flowering time, a trait that has been a subject of many studies on serpentine adaptation. For instance, earlier flowering of serpentine plants was documented in *Solidago virgaurea* (Sakaguchi et al., 2017), *Picris hieracioides* (Sakaguchi et al., 2018), and *Helianthus exilis* (Sambatti and Rice, 2006). Further, selection for flowering time differentiation has been shown in various taxa adapted to serpentine soils in California (Sianta and Kay, 2019). Interestingly, Sianta and Kay (2021) showed that colonization of serpentine sites can cause maladaptive shifts to later flowering in the early stages of divergence. Indeed, this is in congruence with observations in *A. arenosa*, where we observed later flowering in the population S2 relative to N2, i.e., in the population pair exhibiting the most recent divergence time (Konečná et al., 2021). In line with this, the opposite trend for earlier flowering of S3 population relative to N3 when grown on serpentine has been observed in the population pair 3 exhibiting the highest genetic divergence (also in line with earlier observations on this population (Arnold et al., 2016)). Therefore, we hypothesize that genetic divergence may again underlie the observed differences.

### Genetic redundancy and polygenic architecture underlying the observed parallelism

Although significant, gene parallelism affected only a small fraction of candidate genes. This is rather surprising given the large pool of variation shared in this polyploid and recently diverged system (Konečná et al., 2021). However, it is consistent with an array of other studies showing similar levels of parallelism (Lai et al., 2019; Preite et al., 2019; Ji et al., 2020; Bohutínská et al., 2021; James, Wilkinson, et al., 2021; Papadopulos et al., 2021). Although, a possibly high number of false-positive candidates resulting from F_ST_ scans and limits in the detection of soft sweeps (Hoban et al., 2016) can lead to a decrease in the percentage of parallels, another mechanism beyond sharing the same genes in repeated adaptation shall be sought, likely related to genetic redundancy and overall polygenic architecture of serpentine adaptation as it is explained below.

The relatively low number of shared differentiation candidate genes can also reflect a rather simple genetic architecture of serpentine adaptation that is driven by only a few high-effect genes. Although this seems to be indeed the case for several serpentine plants (*Silene, Mimulus;* (Bratteler et al., 2006; Selby and Willis, 2018)), it is likely not the case of *Arabidopsis*. Polygenic basis of serpentine adaptation, i.e. allele frequency shifts in many genes across population pairs (Yeaman, 2015; Wilkinson et al., 2021), has been previously identified by high-density divergence scans for selection in *A. arenosa* and *A. lyrata* (Turner et al., 2010; Arnold et al., 2016; Konečná et al., 2021). These studies showed that many loci with small effects are often under selection when facing complex or even well-defined environmental challenges. In contrast, evidence for simple genetic architecture is typically based on quantitative trait loci (QTL) studies, such as Ni tolerance in *Silene vulgaris* and *Caulanthus amplexicaulis* (Bratteler et al., 2006; Burrell et al., 2012) and survival of *Mimulus guttatus* (Selby and Willis, 2018). However, the QTL studies are usually biased to detect major-effect loci thus providing limited insights into polygenic adaptations.

In line with the hypothesis of genetic redundancy, we also observed similarities in functional pathways, many of which were corresponding to observed parallel phenotypic differences in iron ion homeostasis, inorganic cation transmembrane transport, steroid biosynthetic process; molecular functions of calcium ion binding; and KEGG pathway (Kanehisa and Goto, 2000) of plant hormone signal transduction (Fig. 3C and Fig. 4 for other functional pathways). Importantly, significant overlaps in the GO terms remained even when parallel gene candidates were excluded, demonstrating selection on different genes from similar functional pathways and developmental processes. This shows the important role of genetic redundancy in rapid adaptations with a polygenic basis (Boyle et al., 2017; Barghi et al., 2020; Láruson et al., 2020). That genetic redundancy is an important factor underlying parallel adaptation towards complex environmental challenges that have been also shown in other systems such as in alpine *Heliosperma pusillum* (Trucchi et al., 2017; Szukala et al., 2021) or dune *Senecio lauttus* (James, Allsopp, et al., 2021), but also from more narrowly defined environments – metal-polluted mines in *Silene uniflora* (Papadopulos et al., 2021) or temperature adaptations in *Drosophila simulans* (Barghi et al., 2019). In summary, given the expected polygenic basis of adaptation, it is unlikely that limited gene-level parallelism simply reflects genetic architecture but rather represents an effect of genetic redundancy.

## Conclusions

Here, we document an overall parallel response of plants to serpentine substrate stress across a diversity of functional traits with utmost very limited trade-offs. Repeatedly identified enrichment of multiple relevant functional pathways, despite rather modest gene-level parallelism, demonstrates that a complex interplay of allele sharing, likely polygenic architecture of the adaptation, and genetic redundancy underlies the observed parallel phenotypic manifestation of serpentine adaptation. Despite this general trend, there are also certain population- and trait-specific non-parallel deviations that may reflect neutral processes and/or site-specific selection. Further investigations of the genotype-phenotype links in well-defined strongly selective environments, such as serpentines, will help us to better assess the role of genetic redundancy in polygenic systems and its impact on the extent of parallelism in adaptation and thus evolutionary predictability.

## Author contributions

V.K. and F.K. conceived the study. V.K., F.K., M.C., E.T., and M.S. designed experiments. V.K. and M.C. did field collections. A.K. performed the laboratory analyses. V.K., D.P. and M.S. performed the experiments. V.K. and D.P. performed data analyses. V.K. and F.K. wrote the manuscript with input from all authors.

## Acknowledgements

This work was supported by Charles University (project Primus/SCI/35 to F.K. and the Grant Agency of Charles University project No. 410120 to V.K.) and the European Research Council (ERC) under the European Union’s Horizon 2020 research and innovation programme [ERC-StG 850852 DOUBLE ADAPT to F.K.]. Additional support was provided by the long-term research development project No. RVO 67985939 of the Czech Academy of Sciences and by Charles University Research Centre program No. 204069. Computational resources were provided by the CESNET LM2015042 and the CERIT Scientific Cloud LM2015085. The authors thank Petr Knotek, Tereza Holcová, Veronika Vlčková, Eliška Konečná, Barbora Lepková, Timothée Lamotte, and Karolína Havlíková for help with field work, experiments, data collection and laboratory work.

## Data accessibility statement

Sequence data are freely available under BioProject PRJNA667586 at the https://www.ncbi.nlm.nih.gov/bioproject/PRJNA667586.

