## Supplementary material for "Genomic basis and phenotypic manifestation of (non-)parallel serpentine adaptation in *Arabidopsis arenosa*": Supplemetary Methods, Figures and Tables

### SUPPLEMENTARY METHODS

*Reciprocal transplant experiment*

We germinated seeds from 12 mother plants/population, each representing a seed family of a mixture of full- and half-sibs. Altogether 15 seeds/family/treatment were put in Petri dishes filled by either type of soil. Seeds germinated in a growth chamber (Conviron) under conditions approximating spring season at the original sites: 12 h dark at 10 °C and 12 h light at 20 °C. The germination rates were published in Konečná et al. (2021). Due to germination failure of N1 seeds in S1 soil, we measured the differential growth response on plants that were germinated in non-serpentine soils and were subjected to the serpentine soil treatment later, at a seedling stage. We chose 44–50 seedlings equally representing the progeny of 11 maternal plants per each population (284 seedlings in total), and transferred each individual to a separate pot filled with ~1 L of either its native soil or the alternative paired soil (i.e. S1 soil for N1 population and vice versa). We randomly swapped the position of each pot twice a week and watered the plants with tap water when needed.

*Root growth experiments in Ca + Mg solutions*

The control medium (Ca/Mg ratio 1.97) was based on one-fifth strength Murashige-Skoog medium and contained: 3.76 mM $KNO_3$, 0.25 mM $KH_2PO_4$, 0.3 mM $MgSO_4$, 0.59 mM $CaCl_2$, 20 μM $H_3BO_3$, 0.02 μM $CoCl_2$, 0.02 μM $CuSO_4$, 20 μM $FeSO_4$, 22.4 μM $MnSO_4$, 0.21 μM $Na_2MoO_4$, 1 μM KI, and 5.98 μM $ZnSO_4$. Medium with the Ca/Mg ratio of 0.04 contained 0.2 mM $CaCl_2$ and 4.5 mM of $MgCl_2$ according to Bradshaw (2005). Medium with the Ca/Mg ratio of 0.2 contained 2.15 mM $CaCl_2$ and 6.97 mM $MgCl_2$. Other salts were added in the same concentrations as in the control medium. All media were supplemented with 1 % w/v sucrose and solidified with 1 % w/v agar (Plant agar; Duchefa, Netherlands). The cultivation media were adjusted to pH = 6.0 by adding NaOH.

Seeds were surface sterilized with a 20 % solution of commercial bleach mixed with 0.1 % Triton for 15 min and then washed with distilled water three times. Seeds were sown out on a control medium in 120 × 120 mm sterile plates and cold stratified at 4 °C in dark conditions for 7 days. After the stratification, seeds germinated in a cultivation room under constant growth conditions (22/18 °C day/night temperature, 16/8 hours light/dark cycle). Six days after germination the seedlings were transferred onto media with the Ca/Mg ratios of 0.2, 0.04, and 1.97 and cultivated for the next nine days in the same cultivation conditions. The experiment was terminated 15 days after germination, plants were fixed overnight in 4 % formaldehyde solution, degassed under vacuum, and gradually saturated with 15 % and 30 % glycerol. Roots of the glycerol-saturated plants were scanned at high resolution (1200dpi, 24-bit) and root system traits were measured using Root analyser plug-in in NIS Elements AR 3.22.05 software (Laboratory Imaging).

*Elemental analysis of soil and leaf samples*

The detection and quantification of Ca, Co, Cr, Mg, and Ni in plant tissues of *A. arenosa* and soil samples was carried out using inductively coupled plasma optical emission spectrometry (ICP OES).

Due to very small amounts of leaf samples taken (tens of milligrams), that minimized the effect on fitness of sampled individuals, the samples were decomposed prior to the analysis using a microwave oven Speedwave ®Xpert (Berghof, Germany; maximal applied power 2000 W) with a multi-tube system. The decomposed plant tissue (2 replicates, 8 – 50 mg according to the available sample amount) was inserted into digestion tubes and treated with 2 mL of sub-boilingly distilled nitric acid (Lachner, Czech Republic) under the following conditions: 10 min hold on 170 °C, 30 % of maximal power, 10 min on 200 °C, 30 % of power, 30 min on 30 °C, 0 % of power. The mineralised samples were filled up to the final volume of 10 mL with deionised water (conductivity 0.055 $\mu S.cm^{-1}$, Evoqua Water Technologies, Germany).

The elemental analysis of Ca, Co, Cr, Mg, and Ni was carried out using the sequential, radially viewed ICP OES spectrometer INTEGRA 6000 (GBC, Dandenong Australia) equipped with the ultrasonic nebulizer U5000AT+ (Teledyne Cetac Technologies, the USA), concentric nebulizer (2 $ml.min^{-1}$) and a glass cyclonic spray chamber (both Glass Expansion, Australia). The analytical lines used were Mg 285.2213 nm, Ca 422.673 nm, Ni 221.647 nm, Co 238.892 nm, and Cr 267.716 nm. The operation conditions of the ICP OES analysis were as follows: sample flow rate 1.5 $mL.min^{-1}$, plasma power 1000 W; plasma, auxiliary and nebulizer gas flow rates 10, 0.4, and 0.52 $L.min^{-1}$, respectively, photomultiplier voltage 600 V for Co, Cr, and Ni and 350 V for Ca and Mg, view height 6.5mm, three replicated reading on-peak 1 s, fixed point background correction. The multielement standards containing 10 – 5 – 1 – 0.5 – 0.1 $mg.L^{-1}$ of Mg and Ca and 0.1 – 0.05 – 0.01 – 0.005 – 0.001 $mg.L^{-1}$ of Co, Cr, and Ni were used for instrument calibration. The external calibration standards were prepared using standard solutions of Ca, Co, Cr, Mg, and Ni all containing 1 $g.L^{-1}$ (SCP, Canada). The limits of detection (concentration equal to three times the standard deviation at the point of the background correction) were 0.0005 $\mu g.L^{-1}$ for Co, Cr, and Ni and 2 $\mu g.L^{-1}$ for Ca and Mg. Certified reference material (Bush twigs and leaves GBW 07602 from the China National Analysis Centre for Iron and Steel, Beijing) was used to validate the method and for the quality control.

For the elemental soil analysis, the modified method according the Czech legislation using nitric acid (Zbíral et al., 2016) was used. 10 g fine-grained soil (2 mm) together with 100 mL of the 2 M nitric acid (produced by Lachner, the Czech Republic and sub boilingly distilled as above mentioned) was left standing 16 hours, then shaken on the rotary shaker for 1 hour, filtered and analysed using ICP OES. The elemental analysis of Ca, Co, Cr, Mg, and Ni was carried out using the ICP OES spectrometer operated under the same operational conditions as plants above.

SUPPLEMENTARY TABLES AND FIGURES

**STable 1.** Details on sampled populations.

| **Pop** | **Ploidy** | **Population name** | **Natural populations** | **Experiment** | **Bedrock type** | **Altitude** | **Latitude** | **Longitude** | **Country** |
|---|---|---|---|---|---|---|---|---|---|
| | | | N ind genome/soil ionomics | N ind cultivated in total/ionomics S treatment | | | | | |
| S1 | 4x | Borovsko | 7/8 | 47/10 | serpentine | 416 | 49.68381 | 15.13326 | CZ |
| S2 | 4x | Steinegg | 7/8 | 50/10 | serpentine | 414 | 48.62993 | 15.54256 | AT |
| S3 | 4x | Gulsen | 8/8 | 45/10 | serpentine | 628 | 47.28167 | 14.92764 | AT |
| N1 | 4x | Vlastejovice | 8/8 | 48/7 | siliceous | 345 | 49.73496 | 15.17484 | CZ |
| N2 | 4x | Fuglau | 8/8 | 50/9 | siliceous | 436 | 48.63149 | 15.55723 | AT |
| N3 | 4x | Ingeringgraben | 8/8 | 44/10 | siliceous | 950 | 47.28405 | 14.68154 | AT |

**STable 2.** Overview of survival and transition to reproduction of number of individuals in reciprocal transplant experiment.

| **Pop** | **Treatment** | **Total N of plants** | **Flowering buds** | **Flowers** | **Fruits** |
|---|---|---|---|---|---|
| N1 | S | 24 | 21 | 13 | 6 |
| N1 | N | 24 | 24 | 22 | 20 |
| N2 | S | 25 | 24 | 8 | 3 |
| N2 | N | 25 | 24 | 19 | 14 |
| N3 | S | 22 | 18 | 13 | 8 |
| N3 | N | 22 | 19 | 17 | 15 |
| S1 | S | 23 | 23 | 21 | 17 |
| S1 | N | 24 | 23 | 22 | 21 |
| S2 | S | 25 | 23 | 9 | 4 |
| S2 | N | 25 | 25 | 22 | 15 |
| S3 | S | 23 | 22 | 22 | 22 |
| S3 | N | 22 | 22 | 19 | 17 |

**STable 3.** The effects of substrate of origin, population pair and their interaction on total fitness inferred by an aster hierarchical model. The aster models consisted of four fitness components, which were aligned in the following directional graph: proportion to bolting → proportion to flowering → fruit production → total seed mass production. All factors were tested by likelihood ratio tests using nested null models (model separately with substrate of origin or treatment effects were compared to model including both factors, further the model with both substrate of origin and population pair effects were compared to mode with substrate of origin*population pair interaction).

| **Treatment** | **Tested factor** | **Null df** | **Alternative df** | **Null deviance** | **Alternative deviance** | **Test df** | **Test deviance** | ***p* value** |
|---|---|---|---|---|---|---|---|---|
| serpentine | Substrate of origin | 5 | 7 | 20002 | 20346 | 2 | -344 | < 0.0001 |
| | Population pair | 6 | 7 | 20031 | 20346 | 1 | -315 | < 0.0001 |
| | Substrate of origin + Population pair | 7 | 9 | 20346 | 20384 | 2 | -38 | < 0.0001 |
| non-serpentine | Substrate of origin | 5 | 7 | 28718 | 29566 | 2 | -848 | < 0.0001 |
| | Population pair | 6 | 7 | 29554 | 29566 | 1 | -12 | 0.0005 |
| | Substrate of origin + Population pair | 7 | 9 | 29566 | 29647 | 2 | -81 | < 0.0001 |

**STable 4.** Differences in total fitness inferred by hierarchical aster models separately for each population pair in serpentine and non-serpentine treatment. The differences were tested using likelihood ratio tests.

| **Population pair** | **Serpentine treatment** | | **Non-serpentine treatment** | |
|---|---|---|---|---|
| | **$\chi2$** | ***p* value** | **$\chi2$** | ***p* value** |
| S1-N1 | 19.52 | < 0.0001 | 0.0006 | 0.9798 |
| S2-N2 | 0.45 | 0.5002 | 1.14 | 0.2851 |
| S3-N3 | 366.48 | < 0.0001 | 2.35 | 0.1251 |

**STable 5** Contribution of phenotypic traits to cumulative fitness of *A. arenosa* serpentine and non-serpentine plants.

| Pop | Treatment | N indiv total/total seed mass production/above-ground biomass | Flower production and early survival | Fruit production | Late survival | Total seed mass production | Above-ground biomass | Cumulative fitness |
|---|---|---|---|---|---|---|---|---|
| N1 | S | 24/6/10 | 0.541667 | 0.461538 | 0.833333 | 0.02911 | 0.018178 | 0.00011 |
| N2 | S | 25/3/16 | 0.32 | 0.375 | 1 | 0.103093 | 0.259713 | 0.003213 |
| N3 | S | 22/8/18 | 0.590909 | 0.615385 | 1 | 1.989026 | 1.115717 | 0.806978 |
| S1 | S | 23/17/17 | 0.913043 | 0.809524 | 0.823529 | 0.777088 | 0.191078 | 0.090382 |
| S2 | S | 25/4/13 | 0.36 | 0.444444 | 0.5 | 0.3585 | 0.35725 | 0.010246 |
| S3 | S | 23/22/22 | 0.956522 | 1 | 0.954545 | 2.818786 | 0.927159 | 2.386207 |
| N1 | N | 24/20/16 | 0.916667 | 0.909091 | 0.7 | 0.857097 | 1.143467 | 0.571703 |
| N2 | N | 25/14/12 | 0.76 | 0.736842 | 0.857143 | 1.21923 | 0.955973 | 0.559464 |
| N3 | N | 22/15/14 | 0.772727 | 0.882353 | 0.8 | 0.893986 | 1.02013 | 0.497445 |
| S1 | N | 24/21/11 | 0.916667 | 0.954545 | 0.47619 | 1.136098 | 0.791321 | 0.374591 |
| S2 | N | 25/15/21 | 0.88 | 0.681818 | 0.933333 | 0.795385 | 1.044027 | 0.465026 |
| S3 | N | 22/17/17 | 0.863636 | 0.894737 | 0.882353 | 1.093541 | 0.983422 | 0.733236 |

**STable 6.** Summary of linear models (LM) testing the effect of substrate of origin, the effect of population pair and their interactions on functional traits (ion uptake, morphological life-history traits, and fitness proxies) of plants cultivated in serpentine soils. Tests of significance for individual fixed effect factors and interaction were conducted by Type III Wald $\chi 2$ tests.

| **Response variable** | **Transformation** | **Substrate of origin** | | | **Pop. pair** | | | **Substrate of origin*pop.pair** | | |
|---|---|---|---|---|---|---|---|---|---|---|
| | | Df | $\chi 2$ | *p* | Df | $\chi 2$ | *p* | Df | $\chi 2$ | *p* |
| Ca [ppm] | log | 1 | 18.1277 | *** | 2 | 9.3029 | *** | 2 | 2.6932 | . |
| Mg [ppm] | log | 1 | 5.4663 | * | 2 | 7.3841 | ** | 2 | 2.6947 | . |
| Ca/Mg | log | 1 | 5.569 | * | 2 | 7.8672 | ** | 2 | 15.0728 | *** |
| Co [ppm] | log | 1 | 10.5126 | ** | 2 | 10.7253 | *** | 2 | 1.1056 | ns |
| Cr [ppm] | log | 1 | 22.911 | *** | 2 | 41.466 | *** | 2 | 11.793 | *** |
| Ni [ppm] | log | 1 | 14.3601 | *** | 2 | 62.4837 | *** | 2 | 4.8947 | * |
| Bolting time [days] | log | 1 | 2.0948 | ns | 2 | 26.8956 | *** | 2 | 12.4542 | *** |
| Flowering time [days] | log | 1 | 0.007 | ns | 2 | 2.8453 | . | 2 | 5.116 | ** |
| Rosette area [$mm^2$] | sqrt | 1 | 122.226 | *** | 2 | 6.6118 | ** | 2 | 9.7281 | *** |
| N of additive rosettes | log | 1 | 38.815 | *** | 2 | 11.1697 | *** | 2 | 4.7695 | * |
| Total seed mass production [mg] | log | 1 | 31.0958 | *** | 2 | 7.204 | ** | 2 | 3.3422 | * |
| Above-ground biomass [mg] | sqrt | 1 | 12.2657 | *** | 2 | 12.3582 | *** | 2 | 4.7489 | * |
| Root biomass [mg] | log | 1 | 13.5719 | ** | 2 | 14.7037 | *** | 2 | 3.3024 | . |

*** $p < 0.001$; ** $p < 0.01$; * $p < 0.05$; · $p < 0.1$

**STable 7.** Effects of substrate of origin (S vs. N), treatment (S vs. N), and their interactions on functional traits tested by linear models (LM), binomial generalized linear models (GLM binomial), and linear mixed effect models (LMM) for each population pair separately. Tests of significance for individual fixed effect factors and interaction were conducted by Type III Wald $\chi2$ tests. The random effects in LMM were cultivation batch and Petri dish for cultivation.

| Response variable | Transformation | Model | Pop.pair | Substrate of origin | | | Treatment | | | Ecotype*treatment | | |
|---|---|---|---|---|---|---|---|---|---|---|---|---|
| | | | | df | $\chi2$ | *p* | df | $\chi2$ | *p* | df | $\chi2$ | *p* |
| Bolting time [days] | log | LM | 1 | 1 | 1.6173 | ns | 1 | 5.3656 | * | 1 | 1.9493 | ns |
| Bolting time [days] | log | LM | 2 | 1 | 5.4496 | * | 1 | 9.5134 | ** | 1 | 0.1474 | ns |
| Bolting time [days] | log | LM | 3 | 1 | 60.53 | *** | 1 | 10.794 | ** | 1 | 12.1929 | *** |
| Flowering time [days] | log | LM | 1 | 1 | 0.0067 | ns | 1 | 4.1831 | * | 1 | 0.0465 | ns |
| Flowering time [days] | log | LM | 2 | 1 | 2.1932 | ns | 1 | 2.1896 | ns | 1 | 6.3390 | * |
| Flowering time [days] | log | LM | 3 | 1 | 17.732 | *** | 1 | 5.15737 | ** | 1 | 5.1469 | * |
| Rosette area [$mm^2$] | sqrt | LM | 1 | 1 | 48.676 | *** | 1 | 251.722 | *** | 1 | 33.82 | *** |
| Rosette area [$mm^2$] | sqrt | LM | 2 | 1 | 42.6912 | *** | 1 | 144.2352 | *** | 1 | 7.9263 | ** |
| Rosette area [$mm^2$] | sqrt | LM | 3 | 1 | 85.345 | *** | 1 | 53.6161 | *** | 1 | 35.877 | *** |
| N of additive rosettes | log | LM | 1 | 1 | 25.1521 | *** | 1 | 66.8079 | *** | 1 | 10.4011 | ** |
| N of additive rosettes | log | LM | 2 | 1 | 13.2123 | *** | 1 | 94.4882 | *** | 1 | 2.2259 | ns |
| N of additive rosettes | log | LM | 3 | 1 | 6.4998 | * | 1 | 3.4469 | * | 1 | 4.1851 | * |
| Probability of fruit production (0/1) | | GLM binomial | 1 | 1 | 10.2295 | ** | 1 | 14.043 | *** | 1 | 2.8821 | . |
| Probability of fruit | | GLM binomial | 2 | 1 | 0.1651 | ns | 1 | 9.2197 | ** | 1 | 0.0287 | ns |

| | | | | | | | | | | | | |
|---|---|---|---|---|---|---|---|---|---|---|---|---|
| production (0/1) | | | | | | | | | | | | |
| Probability of fruit production (0/1) | | GLM binomial | 3 | 1 | 10.7342 | ** | 1 | 4.3036 | * | 1 | 5.9476 | * |
| Total seed mass production [mg] | log | LM | 1 | 1 | 27.7629 | *** | 1 | 27.4466 | *** | 1 | 12.937 | *** |
| Total seed mass production [mg] | log | LM | 2 | 1 | 1.5909 | ns | 1 | 7.0795 | * | 1 | 4.2835 | * |
| Total seed mass production [ mg] | log | LM | 3 | 1 | 6.4173 | * | 1 | 0.0225 | ns | 1 | 3.0056 | . |
| Above-ground biomass [mg] | sqrt | LM | 1 | 1 | 6.7994 | * | 1 | 74.0321 | *** | 1 | 9.7637 | ** |
| Above-ground biomass [mg] | sqrt | LM | 2 | 1 | 1.8036 | . | 1 | 39.1427 | *** | 1 | 0.3055 | ns |
| Above-ground biomass [mg] | sqrt | LM | 3 | 1 | 0.3318 | ns | 1 | 0.0068 | ns | 1 | 0.0252 | ns |
| Root biomass [mg] | log | LM | 1 | 1 | 19.738 | *** | 1 | 77.331 | *** | 1 | 17.991 | *** |
| Root biomass [mg] | log | LM | 2 | 1 | 5.6086 | * | 1 | 25.3902 | *** | 1 | 4.8871 | * |
| Root biomass [mg] | log | LM | 3 | 1 | 0.0142 | ns | 1 | 0.5516 | ns | 1 | 5.0671 | * |
| Probability of late survival (after fruit production) (0/1) | | GLM binomial | 1 | 1 | 6.5438 | * | 1 | 8.0743 | ** | 1 | 2.1933 | ns |
| Probability of late survival | | GLM binomial | 2 | 1 | 0.0848 | ns | 1 | 4.8608 | * | 1 | 0.0621 | ns |

| | | | | | | | | | | | | |
|---|---|---|---|---|---|---|---|---|---|---|---|---|
| (after fruit production) (0/1) | | | | | | | | | | | | |
| Probability of late survival (after fruit production) (0/1) | | GLM binomial | 3 | 1 | 0.0001 | . | 1 | 0.1392 | ns | 1 | 0.0001 | . |
| Total root growth [cm] | log | LMM | 1 | 1 | 126.12 | *** | 2 | 61.19 | *** | 2 | 45.36 | *** |
| Total root growth [cm] | log | LMM | 2 | 1 | 72.507 | *** | 2 | 102.225 | *** | 2 | 44.424 | *** |
| Total root growth [cm] | log | LMM | 3 | 1 | 119.729 | *** | 2 | 79.918 | *** | 2 | 26.266 | *** |
| Main root length [cm] | | LMM | 1 | 1 | 15.2207 | *** | 2 | 26.2520 | *** | 2 | 6.9497 | * |
| Main root length [cm] | | LMM | 2 | 1 | 39.866 | *** | 2 | 43.007 | *** | 2 | 19.634 | *** |
| Main root length [cm] | | LMM | 3 | 1 | 92.897 | *** | 2 | 54.702 | *** | 2 | 23.9 | *** |
| Density of lateral roots (number of lateral roots/main root length) | sqrt | LMM | 1 | 1 | 16.617 | *** | 2 | 26.164 | *** | 2 | 10.292 | ** |
| Density of lateral roots (number of lateral roots/main root length) | sqrt | LMM | 2 | 1 | 22.098 | *** | 2 | 20.394 | *** | 2 | 10.882 | ** |
| Density of lateral roots (number of lateral roots/main root length) | sqrt | LMM | 3 | 1 | 156.261 | *** | 2 | 108.780 | *** | 2 | 54.395 | *** |

*** $p < 0.001$; ** $p < 0.01$; * $p < 0.05$; · $p < 0.1$

**STable 8.** Quantification of parallelism by genes and functions. Percentages of shared differentiation candidate genes and significantly enriched gene ontology terms ($p < 0.05$) in biological processes domain - BP, molecular functions - MF, and cellular components - CC between any two population pairs when applying the classic algorithm (Alexa and Rahnenfuhrer, 2020). We calculated proportion of shared genes or functions as number of shared differentiation candidate genes or gene ontology (GO) terms per each two population pairs divided by the number of remaining "unique" (non-shared) differentiation candidate genes or GO terms for selected population pair combination. Percentages of shared enriched gene ontology terms, which were calculated based on GO enrichment without parallel differentiation candidate genes, are given in brackets.

| **Level** | **S1-N1 – S2-N2** | **S1-N1 – S3-N3** | **S2-N2 – S3-N3** |
|---|---|---|---|
| genes | 2.43 % | 2.46 % | 2.21 % |
| enriched gene ontology terms BP | 9.20 % (5.33 %) | 4.63 % (4.43 %) | 2.06 % (1.19 %) |
| enriched gene ontology terms MF | 6.98 % (1.56 %) | 5.75 % (1.63 %) | 8.77 % (1.73 %) |
| enriched gene ontology terms CC | 9.18 % (7.55 %) | 5.88 % (6.12 %) | 5.81 % (2.22 %) |

**STable 9.** Number of protein interactions inferred by STRING for candidate differentiation genes for each population pair.

| **Population pair** | **N differentiation candidate genes** | **N differentiation candidate genes interacted in total[1]** | **N differentiation candidate genes interacted with genes from at least one other population pair[2]** | **Proportion[1]** | **Proportion[2]** |
|---|---|---|---|---|---|
| S1-N1 | 494 | 358 | 340 | 0.72 | 0.69 |
| S2-N2 | 513 | 357 | 332 | 0.7 | 0.65 |
| S3-N3 | 504 | 363 | 333 | 0.72 | 0.66 |

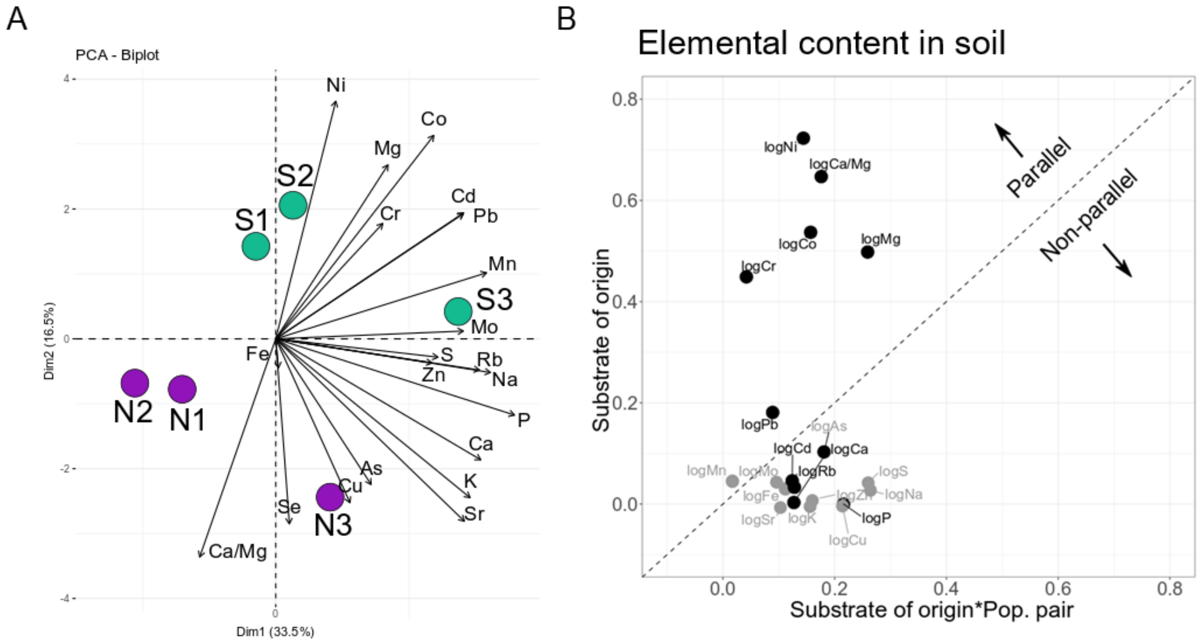


**SFig. 1.** A) PCA based on all soil variables (population means of 7/8 individual observations - STable 1) measured at original sites showing the differentiation between S and N soils. Serpentine (S, green dots) and non-serpentine (N, violet dots). B) Variation in the extent of parallelism among individual soil elemental concentrations from natural populations. The extent of parallelism was estimated as effect sizes (Eta-squared) in linear models addressing the effect of substrate of origin (S vs. N), pop. pair (1, 2, 3) and their interaction, calculated separately for each trait (particular elemental soil concentration and Ca/Mg ratio). The effect size of substrate of origin (y-axis) shows the extent to which a given trait diverges predictably between substrate of origins, i.e., in parallel, while the substrate of origin*pop. pair effect size (x axis) quantifies the extent to which serpentine/non-serpentine soil divergence varies across population pairs (i.e., deviates from parallel). Points falling above the dashed line have a larger substrate of origin effect (i.e. parallel) than substrate of origin*pop. pair interaction effect (i.e. non-parallel). Note: green arrows indicate the trend in all serpentine populations; grey points indicate the non-significant ($p < 0.05$) effect of substrate of origin.

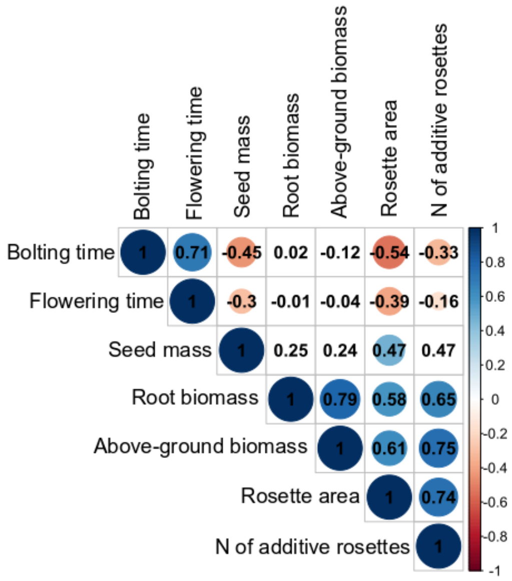


**SFig. 2.** Pairwise Spearman’s correlations among fitness traits. Note: circle size denotes significance (larger circle=lower *p* value), displayed are only circle sizes with $p < 0.05$).

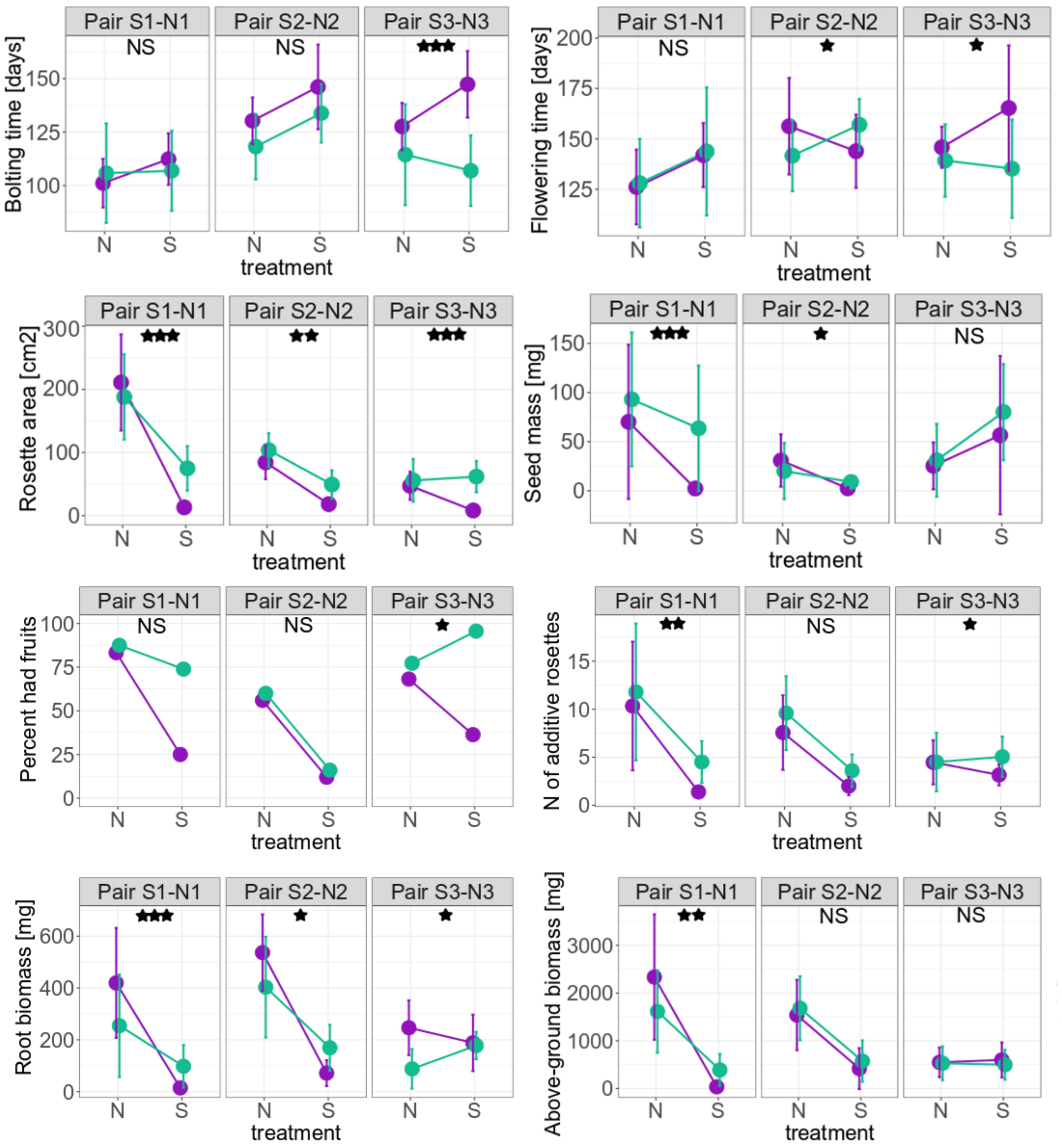


**SFig. 3.** Differences in vegetative and reproductive fitness traits of three population pairs cultivated in local serpentine (S) and non-serpentine (N) soils. Note: asterisks denote the significance of substrate of origin*treatment interaction within each population pair (*** $p < 0.001$; ** $p < 0.01$; * $p < 0.05$; see Table S7); the percentage of plants that had fruits was counted from the total number of cultivated plants.

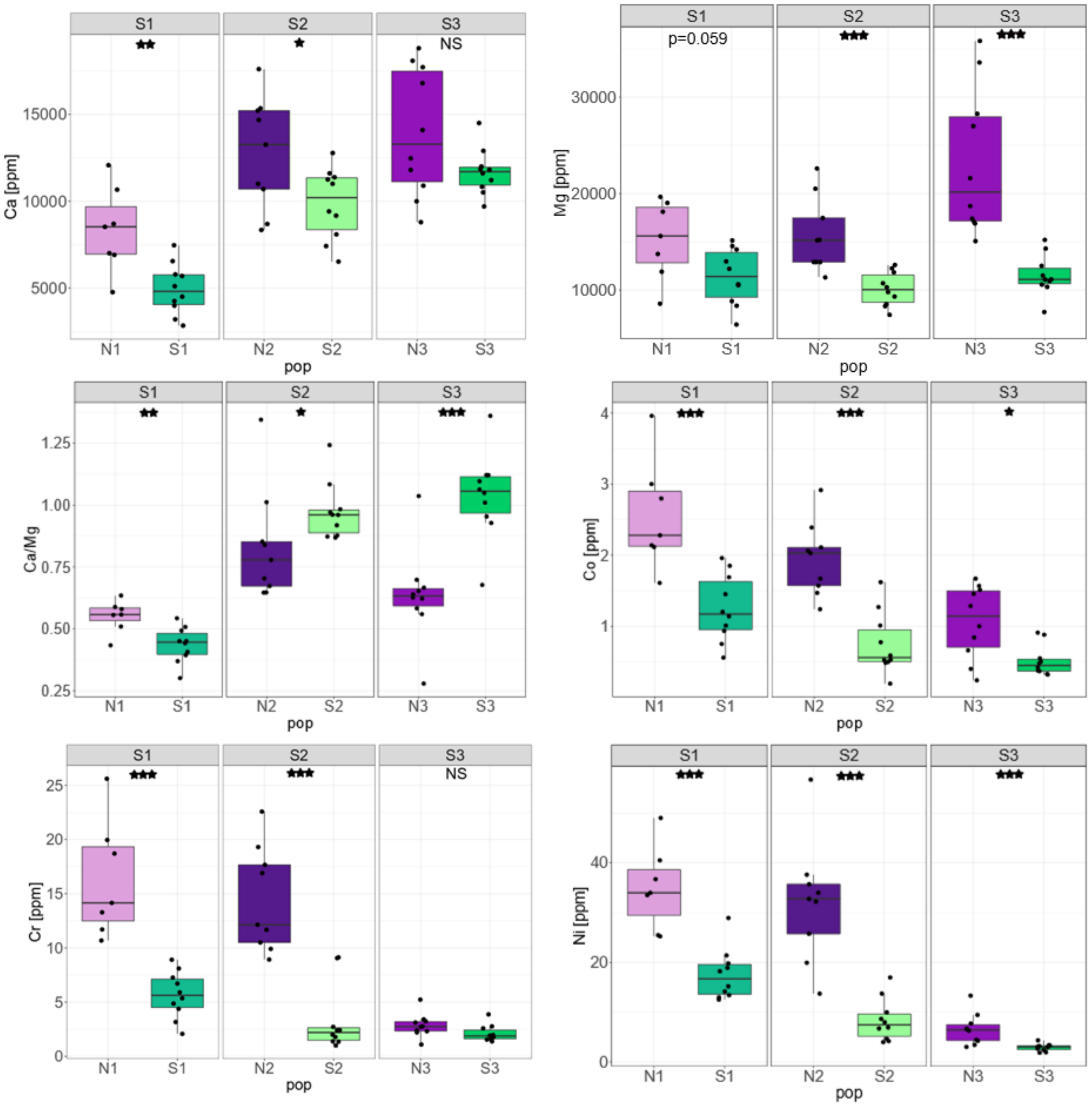


**SFig. 4.** Variation in the uptake of Ca, Mg, and exclusion of Ni, Co, and Cr in serpentine (S) and non-serpentine (N) plants cultivated in serpentine soils (S1, S2, and S3). Note: asterisks denote the significance of the effect of substrate of origin (*** $p < 0.001$; ** $p < 0.01$; * $p <= 0.05$); elemental concentrations were assessed from plants after 2.5 months of cultivation in serpentine soils; dots denote individuals.
